# *subsppLabelR*: a wrapper package in R to automatically label and filter subspecies occurrence data

**DOI:** 10.64898/2026.01.05.697764

**Authors:** Kaiya L. Provost

**Affiliations:** Department of Biology, Adelphi University, One South Avenue, Garden City NY, 11530 USA

**Keywords:** data filtering, occurrence data, species distribution models, subspecies, subsppLabelR

## Abstract

Occurrence data is the basis for many fundamental ecoevolutionary analyses, and many ways of filtering them to be robust enough for analysis have been developed. One issue that still remains is separating out the boundaries of taxa, especially taxa below the species level which often have vague definitions.*subsppLabelR* is an R package that uses labeled data on taxa to automatically define boundaries between them, with various levels of uncertainty. I tested the features of the package on three species of bird that vary in the number and location of subspecies, as well as one sister-species pair with a known overlap in their distribution. I then used existing ecological niche modeling software to compare and contrast their niche space.*subsppLabelR* performs well in scenarios where subspecies are well-sampled and cover large geographic areas, but rare or highly endemic subspecies are difficult to resolve without further input.This package serves to automatically define geographic boundaries of taxa, identifying any sympatric overlaps, and provides an alternate way to clean occurrence data for ecological and evolutionary analyses.

**Data/Code for peer review statement:** Code and data are available with peer review. Raw Data and the package are available as .ZIP files as well as an R script.

## Introduction

Named taxa within species, specifically subspecies, have been controversial in taxonomy and systematics, particularly for birds (e.g., Zink. 2004; Phillimore and Owens, 2006; Winker 2010). IOC 15.1 lists 19,774 subspecies of bird with 2-42 subspecies per each of the 5,004 species with sub-specific taxonomic rankings (Gill et al., 2025). Depending on the group, subspecies can be strictly geographical, reflect known or suspected differences in phenotype, or accurately capture within-species population structure. Multiple lineages exist within bird species, but there is also evidence that avian subspecies do not correspond neatly to these lineages (Zink, 2004; Barrowclough et al., 2016).

Though their utility has been debated (e.g., Meikle, 1957; Mayr et al., 1982; Phillimore and Owens, 2006; Braby et al., 2012), subspecies often serve as shorthand for regional populations. Subspecies are often hypothesized incipient species that may give insight into the process of speciation itself. In birds, *Zonotrichia leucophrys* subspecies are known to have behavioral, morphological, and genetic differentiation (Chilton et al., 2020) but while groups of subspecies in *Cardinalis cardinalis* differ in niche, individual subspecies do not reflect distinct evolutionary entities (Smith et al., 2011; Smith and Klicka, 2013). Even when subspecies represent evolutionary units, many are poorly defined and difficult to demarcate.

Despite subspecific difficulties, they are frequently assigned to individuals. Natural history museums often label specimens to subspecies based on expert opinion or the literature. While (in the author’s experience) miscategorization at the subspecies level is more common than miscategorization at the species level or above, such labels offer insight into where the boundaries actually are. Thus, labels can be leveraged to make inferences about subspecies boundaries, identify putative evolutionary units, and test for differences between those units.

Here, I develop a framework to automatically extract subspecies boundaries from published specimen occurrence databases, *subspplabelR* (pronounced “subspecies labeler”). It is a package implemented in R version 4.4.1 (R Core Team, 2024). I use five taxa to demonstrate its utility, investigate parameter space, and test for differences in climatic niche between subtaxa.

## Materials and Methods

*subspplabelR* downloads occurrence records from available databases, algorithmically labels the records by subspecies, and identifies mislabeled individuals. The labeling procedure integrates existing R packages (Venables and Ripley, 2002; Pebesma and Bivand, 2005; Bivand et al., 2013; Chamberlain, 2017; Bivand and Rundel, 2017; Wickham et al., 2017; Hijmans et al., 2017; Garnier, 2018; Hijmans, 2025). The procedure makes the following assumptions. First, subspecies are distinct units and their boundaries do not overlap. Sympatry is ignored. Second, the vast majority of subspecies labels are correct. Third, the area of densest points for a given subspecies is the area of highest confidence in those subspecies labels. However, sampling bias can lead to poor estimates of where organisms are actually distributed (e.g., Beck et al., 2014, Hughes et al., 2021). Lastly, if subspecies boundaries are already known or approximated using the *subspplabelR* methodology, any occurrences falling within those boundaries can be labeled with confidence.

First, occurrence data are acquired either from user-submission or downloading from occurrence datasets. The latter assumes that you are downloading subspecies within species or species within genera. User-provided data will work for any taxonomic combination (genera within families, etc). To download, a maximum number of points is given that will be downloaded for each taxon from each database. Occurrence points are first downloaded for the full species and are labeled as belonging to an unknown subspecies, even if there are subspecies designations within the data. After that, each subspecies is downloaded from the occurrence databases independently and labeled with the name of that subspecies. Users must ensure that they acquire appropriate DOIs for their data. Downloaded data varies based on the exact databases used. For use in the *subsppLabelR* package, the data must contain the following columns: “name” which gives the full scientific name associated with occurrences, “longitude”, “latitude”, and “subspecies” which gives the initial assumed label. The subspecies column will include the label “unknown”.

After occurrence data are acquired, the package will automatically clean the data with specific steps. Latitudes and longitudes are rounded to the nearest two decimal places to combat uneven geographic sampling. Duplicate localities with the same assumed subspecies are removed. Further filtering can also be done using external packages. Occurrence points are sometimes mislabeled with respect to their coordinates or with respect to their subspecies designation. While manual checking of the data can be done, *subsppLabelR* will perform outlier detection using multivariate Gaussian distribution anomaly detection (Figure 1). This detection is performed for the whole species distribution as well as each labeled subspecies individually. A cutoff value (epsilon) for probabilities is set, and any point with a probability lower than the cutoff is designated an outlier. *subsppLabelR* will allow you to set different cutoffs for the species comparison and the subspecies comparisons.

**Figure 1.**
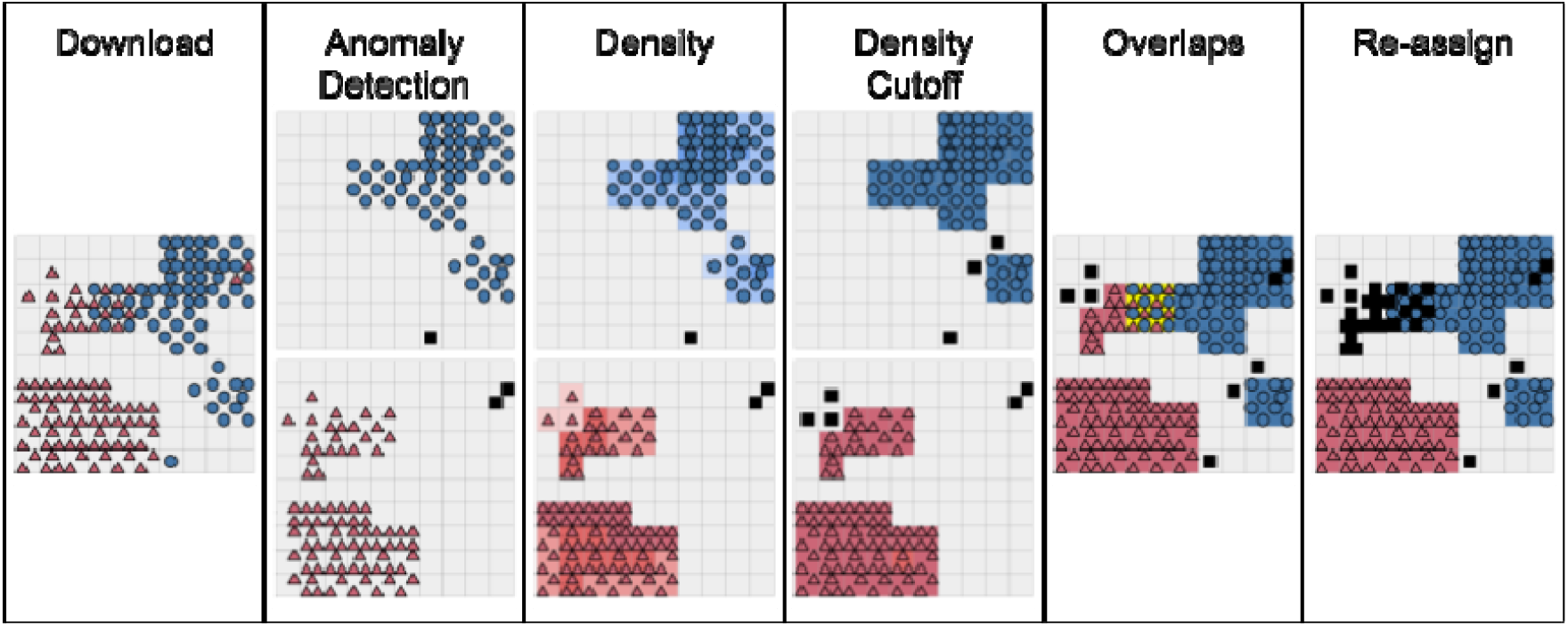
Example workflow for two subspecies. Occurrence data are given as red triangles (subspecies 1), blue circles (subspecies 2), or black squares (unknown). Boxes underneath represent raster pixels. The number of occurrence points within the raster pixel corresponds to color, with lighter shades indicating less density. Yellow boxes overlap.

After occurrence points are cleaned, a density raster is calculated from all points in each subspecies. This is done with two-dimensional kernel density estimation via the kde2d() function in *MASS* version 7.6-60.2 (Venables and Ripley, 2002). The density estimation will not estimate densities of zero, but will instead asymptotically approach zero, even for cells with no points. Cells within the density rasters are then rescaled from 0 to 1, where 0 indicates the least dense cell and 1 indicates the most dense cell.

After this, a user-set percentile cutoff is applied to the data, such that only cells at or above the Nth percentile of the density raster are kept as potential locations for that subspecies. Higher percentile cutoffs will keep smaller and smaller areas. Overlaps between the density rasters are checked and removed for each labeled subspecies (Figure 2). Regions of overlap are preferentially removed from the least dense subspecies range to preserve subspecies whose ranges are very restricted.

**Figure 2.**
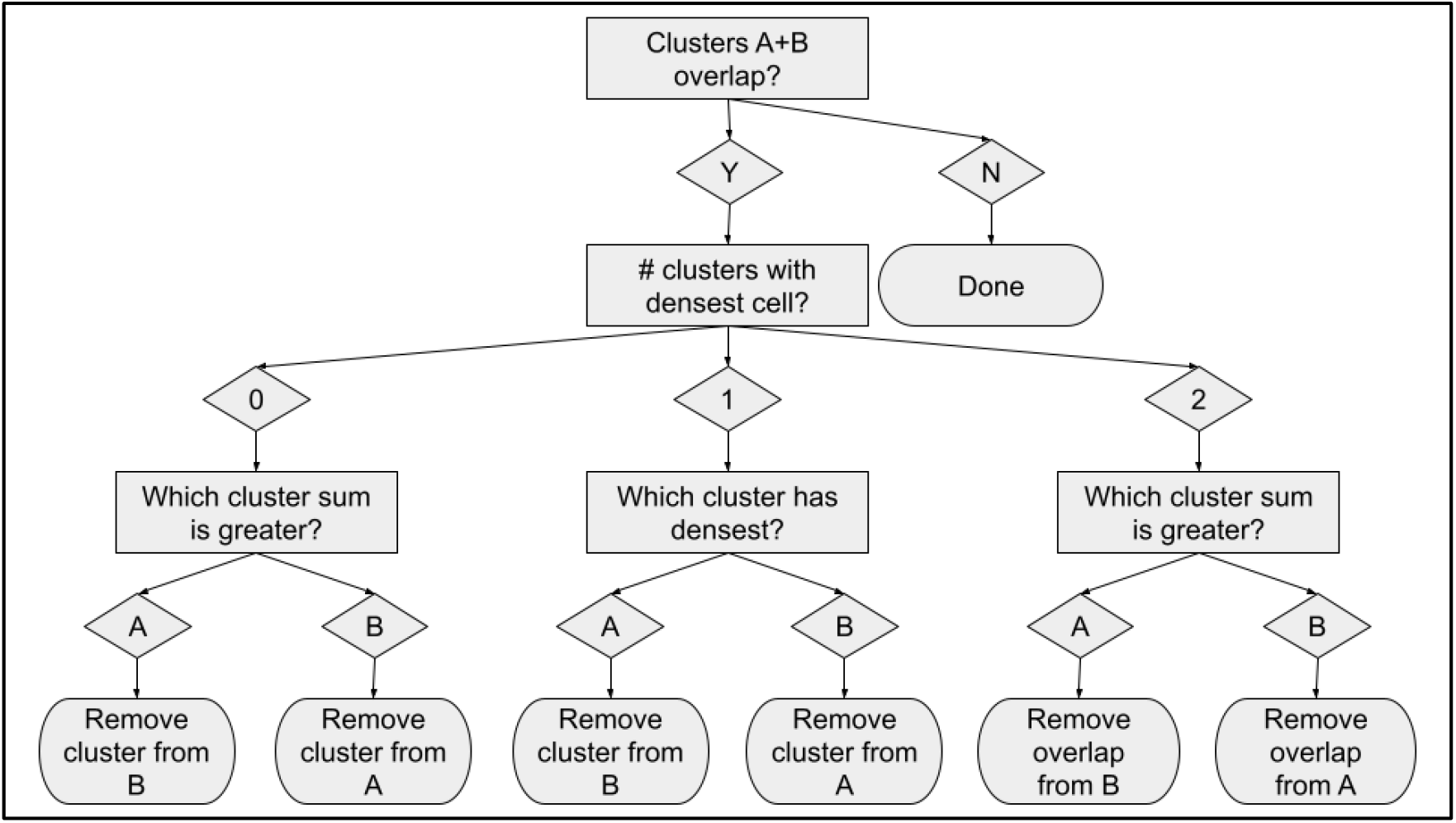
decision tree to determine overlaps between clusters of two different subspecies A and B.

All points, including those that were already labeled, are labeled again based on the non-overlapping density rasters above. If a point falls into one of those rasters, it is given the corresponding label. Points that do not fall into any raster are left as “unknown”. After all points have been assigned, the package flags “suspect” points that will need further human revision. Points are considered suspect if they fall into multiple rasters, have multiple subspecies labels, or have received no labels and are still unknown subspecies. All other points are considered to be confidently assigned.

The methodology outlined above was tested on five North American bird taxa that all reside in deserts. Three species were used to test the separation of subspecies: *Phainopepla nitens* (*nitens* and *lepida*), *Cardinalis sinuatus* (*sinuatus, peninsulae*, and *fulvescens*), and *Toxostoma curvirostre* (*celsum, curvirostre, insularum, maculatum, oberholseri, occidentale*, and *palmeri*). These taxa vary in whether they are diagnosable morphologically, genetically, or geographically (Pyle, 1997; Chu and Walsberg, 2020; Tweit and Thompson, 2020; Provost et al., 2022; Tweit and Thompson, 2020; Rojas-Soto et al., 2007; Zink and Blackwell-Rago, 2000). Of note, *T. c. insularum* is an island endemic and severely range restricted compared to the other six subspecies (Tweit, 2020). An additional test was performed to see if the pipeline worked on two sister species, *Geococcyx californianus* and *G. velox*, which overlap geographically.

Using *subsppLabelR*, I downloaded 2000 maximum occurrences for each subtaxon in the above. A GBIF DOI for these data is pending. After preliminary analysis, I set the epsilon thresholds for outlier detection to be 1x10^-6^ for supertaxa and 1x10^-3.5^ for subtaxa. The higher threshold for subtaxa means that points are more likely to be designated as outliers. Each taxon was tested with the following percentile cutoffs for determining density rasters: 50th, 60th, 70th, 80th, 90th, and 95th (the default). Changing the cutoff thresholds adjusted the amount of overlap, the patchiness in the resulting density rasters, the estimated size of the range, and how many points were successfully labeled. These thresholds were examined by eye and then the “best” threshold was determined to be the one that minimized overlaps and misidentified subspecies while retaining the most range.

To show an example for what subsppLabelR outputs may be used for, I built ecological niche models using the identified occurrence data for *C. sinuatus, P. nitens*, and the *Geococcyx* species. Niche modeling for *T. curvirostre* was excluded due to the large number of pairwise comparisons that would need to be accounted for with this species. I also used rasterized climate data from the WorldClim 2.0 database (Fick and Hijmans, 2017). Niche model selection was performed using *ENMeval* (Muscarella et al., 2014), which optimizes MaxEnt models (Phillips et al., 2006) for different sets of feature classes and regularization values. Final models were then built with MaxEnt and thresholded with *dismo* (Hijmans et al., 2017). Models were built for each subspecies individually. Niches were then compared using *ENMTools* (Warren et al., 2010) specifically to calculate niche overlap, niche equivalency, and niche similarity. A principal components analysis based method (Broennimann et al., 2012) computes the niche space occupied by each species relative to the background niche space available. These values are then used to calculate measures of niche overlap, Schoener’s D and a modified Hellinger’s I, and then finally simulations are used to compare the actual overlap values with null expectations with *ecospat* (Warren et al., 2008; Di Cola et al., 2017). Each simulation was repeated 1000 times to generate 1000 pseudoreplicates. Specifically, these analyses examine whether, compared to random niches, niches are more different than expected.

There was no use of generative AI in any of the work presented herein.

## Results

After comparing the outputs for each of the density percentiles, I selected a cutoff of 0.6 for the *Geococcyx* subtaxa, 0.7 for the *Phainopepla* subtaxa, 0.9 for the *Cardinalis* subtaxa, and 0.95 for the *Toxostoma* subtaxa (Figure 2). Anecdotally, the larger number of subtaxa that are being analyzed, the higher a density cutoff seems to be appropriate. As the density cutoff increases, ranges that are predicted for each subtaxa become smaller and more patchy (Figure 2).

There is a tradeoff between the size of ranges of each subspecies and the amount of overlap between those ranges. Subtaxa are separated by an unrealistically large space with these cutoffs (Figure 3). Generally, the areas that are predicted as each subtaxon are the center of the cloud of points and correspond to the known subspecies descriptions.

**Figure 2.**
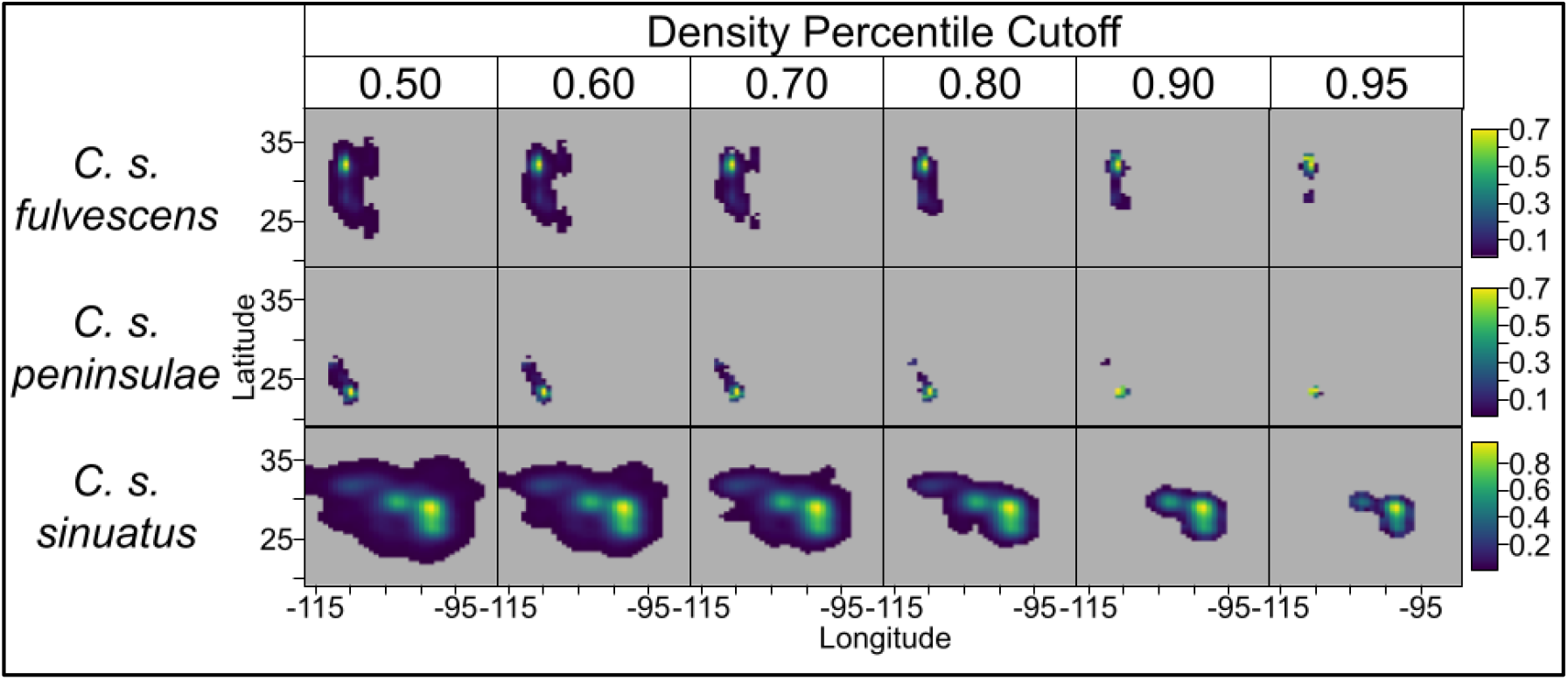
Visualization of density percentiles used for cutoffs of different subspecies of *Cardinalis sinuatus*. Columns show the subspecies range estimated for varying density percentile cutoffs. Rows show the modeled subspecies. X-axis shows longitude. Y-axis shows longitude. Colors indicate densities, with denser regions being warmer colors. Grey indicates the background.

**Figure 3.**
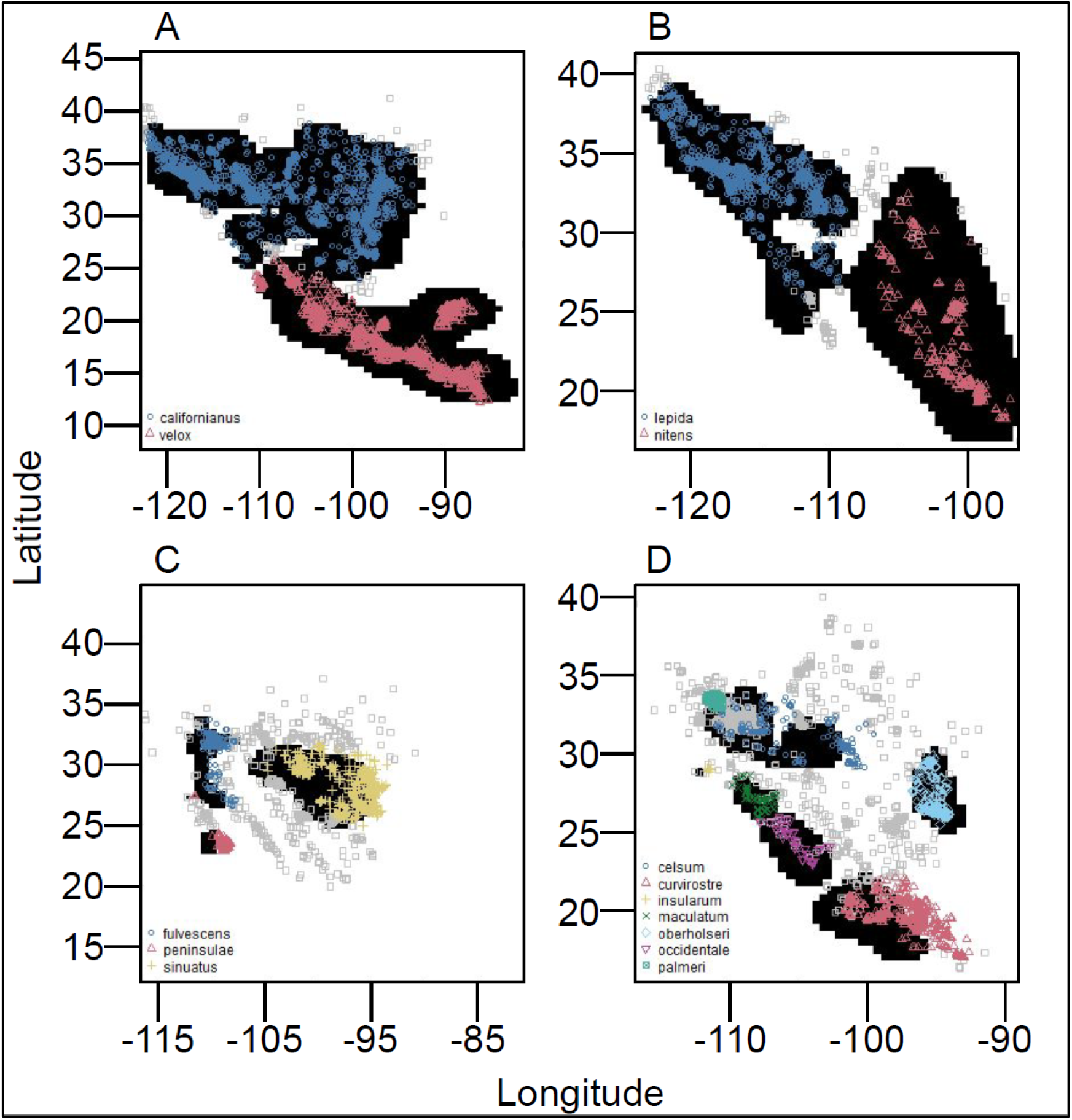
Predicted subspecies identities for four taxa. Y-axis indicates latitude and x-axis indicates longitude. Black squares indicate areas where all subspecies are estimated. Grey squares show points that were removed. A: *Geococcyx*. B: *Phainopepla nitens*. C: *Cardinalis sinuatus*. D: *Toxostoma curvirostre*. Color and shape of points indicate predicted species (A) or subspecies (B, C, D).

Climatic niches were estimated for each subspecies pair within species. In all cases, Hellinger’s I estimates of overlap were higher than Schoener’s D overlaps (Table 1). For *C. sinuatus*, all three subspecies pairs show that niches are less equal than expected (p ≤ 0.03; Table 1). *Cardinalis sinuatus fulvescens* is more similar to *C. s. peninsulae* than expected, though the reverse is not true (Table 1). *Phainopepla nitens* subspecies are also more equal than expected (p = 0.002; Table 1). For the two species of *Geococcyx*, Hellinger’s I and Schoener’s D metrics give opposite results for equivalency, with the former showing significantly more equal than expected and the latter showing significantly less equal than expected. Schoener’s D assumes that niche suitability is proportional to abundance, while Hellinger’s I makes no such assumptions (Warren et al., 2008; Warren et al., 2010).

**Table 1.** Niche metrics for *Cardinalis, Phainopepla*, and *Geococcyx* taxa. Niche overlap, niche equivalency, and unidirectional niche similarity between taxa are calculated for Schoener’s D and Hellinger’s I. Significant differences are marked with an “(L)” for lower than expected and an “(H)” for higher than expected. Equivalency and similarity sections show p-values.

| Taxon A | Taxon B | Niche Metric | Schoener's D | Hellinger's I |
| --- | --- | --- | --- | --- |
| *Cardinalis sinuatus fulvescens* | *C. s. peninsulae* | Overlap | 0.169 | 0.347 |
|  |  | Equivalency | **0.030 (L)** | **0.006 (L)** |
|  |  | Similarity A to B | **0.032 (H)** | **0.020 (H)** |
|  |  | Similarity B to A | 0.360 | 0.384 |
| *C. s. peninsulae* | *C. s. sinuatus* | Overlap | 0.016 | 0.037 |
|  |  | Equivalency | **0.012 (L)** | **0.002 (L)** |
|  |  | Similarity A to B | 0.142 | 0.140 |
|  |  | Similarity B to A | 0.194 | 0.519 |
| *C. s. fulvescens* | *C. s. sinuatus* | Overlap | 0.213 | 0.381 |
|  |  | Equivalency | **0.002 (L)** | **0.002 (L)** |
|  |  | Similarity A to B | 0.913 | 0.741 |
|  |  | Similarity B to A | 0.490 | 0.455 |
| *Phainopepla nitens lepida* | *P. n. nitens* | Overlap | 0.381 | 0.559 |
|  |  | Equivalency | **0.002 (H)** | **0.002 (H)** |
|  |  | Similarity A to B | 0.100 | 0.116 |
|  |  | Similarity B to A | 0.288 | 0.323 |
| *Geococcyx californianus* | *G. velox* | Overlap | 0.143 | 0.210 |
|  |  | Equivalency | **0.002 (H)** | **0.002 (L)** |
|  |  | Similarity A to B | 0.845 | 0.927 |
|  |  | Similarity B to A | 0.533 | 0.561 |

## Discussion

The package *subsppLabelR* is capable of outlining, with various degrees of certainty, the locations of subtaxa within species or genera. There are considerations for end users to address before using the package for their own purposes. Nominate subspecies tend to be over-represented in datasets. Anecdotally, the author has noticed when volunteer workers are digitizing specimens into databases, they tend to put the nominate as a subspecies even when no subspecies is given. This could explain why specimens may be erroneously identified incorrectly. If using only *subsppLabelR* to address this issue, I would recommend exploring the settings of the density cutoffs.

Species and subspecies distributions can actually overlap, despite the package assuming that they do not. Sympatric species, in particular, will have real overlaps removed (e.g., *Geococcyx*; Soberanes-González et al., 2020; Hughes, 2020). Range-restricted subtaxa can also cause problems for the package, especially when being estimated at the same time as more widespread subspecies. For example, *T. c. insularum*, being an island endemic, is estimated to occur in only a single pixel of the underlying species distribution (Tweit, 2020). Adjusting the density cutoffs to retain all taxa may be necessary for the investigator.

In a similar vein, the package at present does not fully resolve sampling bias as it uses a density-based approach. While duplicate latitude-longitude pairs are removed, pixel sizes are large (defaulting to 100x100 pixels for the entire study area, which in this case covers ∼40° latitude and ∼35° longitude). Highly-sampled areas are likely to present as dense. Investigators wishing to adjust for sampling bias may wish to perform additional spatial thinning (such as with *spThin*, Aiello-Lammens et al., 2015) or otherwise adjust for sampling bias (e.g., Hughes et al., 2021). Taxa that experience seasonal migration will also need to be considered. For example, *Phainopepla nitens* is a species with both short-distance migrants, long-distance migrants, and elevational migrants (Chu and Walsberg, 2020). Researchers working on migratory species may wish to subset their points to only the breeding ranges.

In conclusion, this study finds that subspecies boundaries can be automatically extracted using published specimen data. Individuals using this new package *subsppLabelR* should take into consideration multiple properties of their focal taxa, such as range size, susceptibility to sampling bias, seasonal migrations, and sympatry. This package can also be used with other downstream applications, for example estimating niche differentiation between populations.

## Acknowledgements

I would like to thank Rob Harbert, Mary Blair, Peter Galante, and the late Eleanor Sterling for instrumental help developing this project. Helpful feedback was also provided by the New York Regional Species Distribution Modeling Discussion. I would also like to acknowledge all individuals who have contributed to the species occurrence record databases that underlie this work.

The authors declare no conflicts of interest.

